# Learning to Attend Through Value-Based Hypothesis Testing

**DOI:** 10.64898/2026.03.11.710688

**Authors:** C. Maher, I. Saez, A. Radulescu

**Affiliations:** Center for Computational Psychiatry, Icahn School of Medicine at Mount Sinai; Department of Neuroscience, Icahn School of Medicine at Mount Sinai; Department of Neurosurgery, Icahn School of Medicine at Mount Sinai; Center for Advanced Circuit Therapeutics, Icahn School of Medicine at Mount Sinai

## Abstract

In complex environments, humans must determine which features are relevant for learning and decision-making. Psychological theories offer competing accounts of this process: associative models suggest that attention emerges gradually through learned changes in feature values, whereas hypothesis-driven accounts propose that learners selectively attend to actively tested rules. Because attentional states are covert, similar behavior can arise from different underlying strategies, making these accounts difficult to distinguish using choice data alone. We inferred latent attention dynamics during learning and decision-making by training recurrent neural networks on synthetic data generated from feature-based reinforcement learning (FRL) and serial hypothesis testing (SHT) models. A network trained on hybrid (FRL+SHT) data outperformed single-model networks, decoding latent human attention with more than 80% accuracy. These results suggest that human attention reflects an interaction between value-based learning and hypothesis testing, in which learned feature value guides the generation and evaluation of candidate rules.

## Introduction

Natural environments are high-dimensional and noisy, requiring individuals to determine which features are relevant for learning and choice. Selective attention plays a key role in this process by guiding which stimulus features are prioritized for learning and decision-making^1,2^. When choosing a restaurant for instance, agents must infer relevant features (e.g., cuisine, price) from ambiguous feedback. However, there is no unified theoretical account of how attentional selection emerges during learning, with competing frameworks positing fundamentally different underlying mechanisms. Moreover, identical choices can arise from distinct internal processes, rendering latent attention fundamentally non-identifiable from behavior alone.

Two major classes of computational cognitive models, each supported by choice data and neural evidence^3–10^, offer competing accounts of how attention learning is implemented in such circumstances: feature-based reinforcement learning (FRL), in which attention emerges gradually through retrospective value updating of stimulus features^4,11–15^, and serial hypothesis testing (SHT), in which attention is allocated prospectively by sampling and evaluating discrete hypotheses about task-relevant features^5,16,17^. Hybrid models incorporating elements of both mechanisms have also been proposed^6^ and found to best capture human choice behavior in a multidimensional RL task^6^. Importantly, standard FRL accounts rely on slow, incremental updating and struggle to explain the rapid formation and reconfiguration of internal task representations observed in human behavior^5^, suggesting that additional mechanisms may be required to explain attention learning in service of state representation during real-world, multidimensional RL^8,18^.

Model fitting to choice data shows that both FRL and SHT can produce similar choice trajectories in multidimensional RL tasks^5,8^. That is, competing mechanisms can generate nearly indistinguishable observable behavior. However, successful reconstruction of choice does not necessarily reveal the cognitive process that drove the decision. For example, a model may correctly predict that a participant chose a particular restaurant yet fail to reveal which feature of the restaurant guided that choice. As a result, fitting models to choice data alone cannot adjudicate between competing mechanisms underlying attention learning.

To address this, we leveraged LaseNet^19^, a novel RNN-based method for direct inference of latent cognitive variables. We first simulated synthetic behavior during a multidimensional RL task (Figure 1A) under competing cognitive models of attention learning (Figure 1B), allowing us to generate trial-by-trial attentional states with known ground truth. Our models included a Hybrid model that integrates value learning and hypothesis testing^6^. We then trained decoders to recover these latent attentional states from observable choice behavior. If a decoder trained under a given generative cognitive mechanism does capture the structure of attention learning in humans, it should decode human attentional states with high accuracy from human choice data in which attention is independently measured. After training LaseNet on model-specific synthetic data (Figure 1C), we evaluated how well each trained network decoded latent attentional states in a human dataset in which attention was measured via trial-by-trial self-report (Figure 1D).

**Figure 1.**
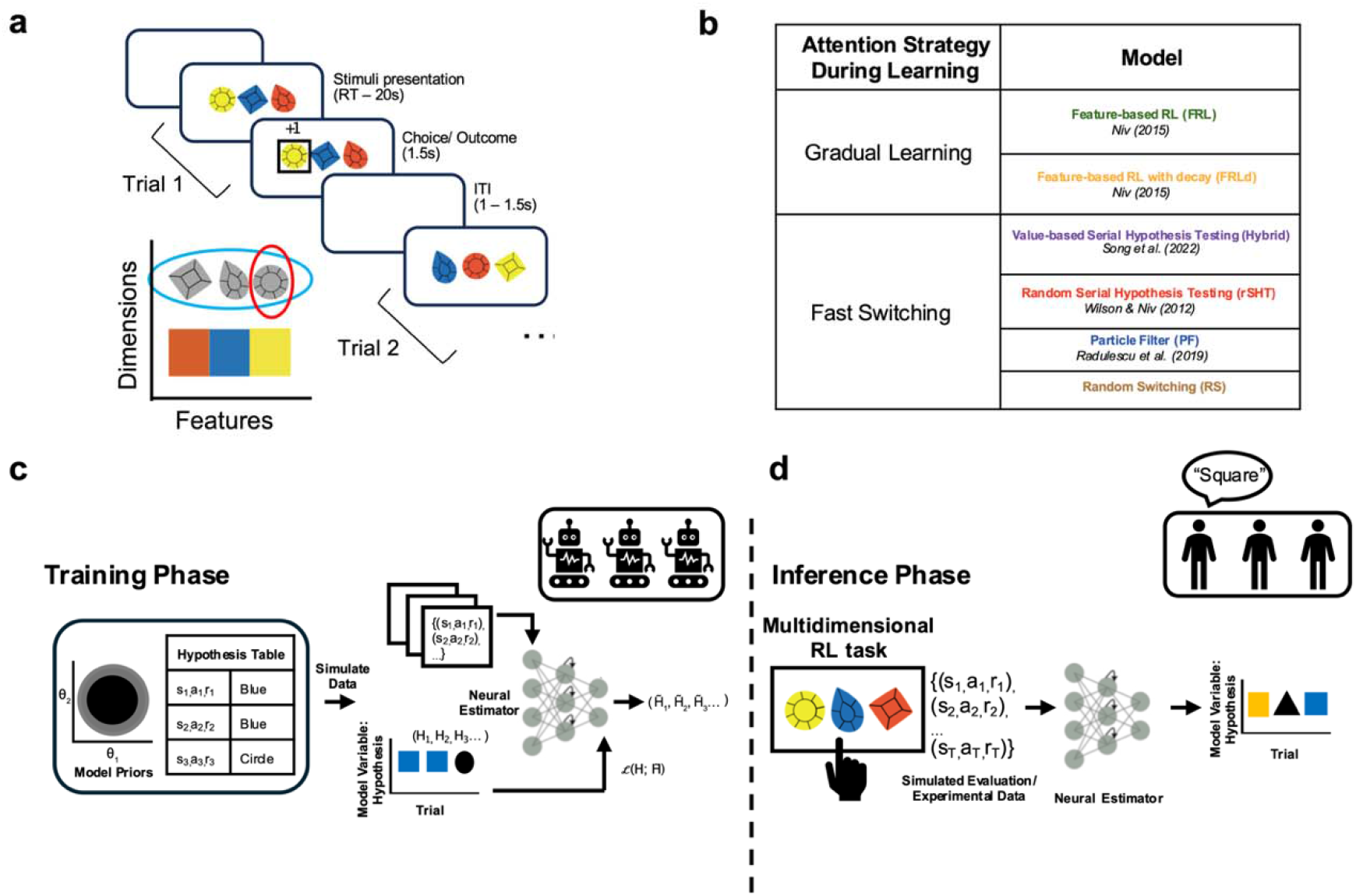
Materials and methods. **A**. Multidimensional RL task. On each block (6 blocks total; 18 trials per block), participants learned via trial and error which dimension was relevant (i.e., “shape”) and within it which feature was most likely to result in reward (i.e., “circle”; 80/20 reward probability). **B**. Overview of the cognitive models evaluated in this study. Models are grouped according to their assumptions about how attention learning evolves: via gradual, continuous updating of feature values (FRL class) versus rapid, discrete hypothesis switching (SHT class). **C**. Network is trained to predict latent variables from a cognitive model (i.e., attended feature) using simulated data. Input includes trial-wise observable data (stimuli, actions, rewards). **D**. Trained networks predict latent variable for experimental data. Schematic adapted from Pan et al.^19^.

We hypothesized that networks trained on synthetic data generated by distinct cognitive mechanisms would acquire mechanism-specific inductive biases^20^. Specifically, we predicted that (1) networks trained under one strategy would generalize poorly to data generated by another, indicating limited cross-mechanism generalization, and (2) a network trained on Hybrid (FRL + SHT) data would most accurately decode trial-by-trial attention in human behavior.

We found that all networks successfully decoded held-out synthetic data generated by their own training model, confirming within-model recovery and validating the analytical framework. Additionally, all networks showed poor cross-model generalization, indicating that each acquired mechanism-specific inductive biases rather than serving as general-purpose decoders. Finally, the Hybrid (FRL + SHT) network most accurately decoded human attentional states, outperforming networks trained on single-mechanism models. Together, these findings support a hybrid account in which human attention learning reflects hypothesis testing complemented by feature-based value learning.

## Results

### Distinct cognitive models propose competing generative mechanisms for attention learning under partial observability

Different computational models propose distinct mechanisms for attention learning during multidimensional reinforcement learning (RL). We investigated attention learning using “Gem Hunters”, a variant of the Dimensions Task^4,5,8–10^ (Figure 1A; see Methods). Briefly, Gem Hunters is a three-armed, multi-round probabilistic reward paradigm in which participants are presented three stimuli (‘gems’) that vary along two visual dimensions, shape and color. Choosing stimuli containing a ‘target feature’ (e.g. ‘circle’) from one of the two dimensions (‘shape’) is rewarded with higher probability than choosing stimuli that do not. The player’s goal is to learn which feature within the relevant dimension maximizes reward. This design allowed us to isolate how selective attention and value learning interact to maintain efficient state representations when the relevant dimension is known. In such environments, attention is thought to support state representation learning by filtering irrelevant information^2,21^, thereby rendering learning tractable and mitigating the curse of dimensionality^22^.

We simulated behavior under two classes of cognitive models, each of which formalizes a different mechanism by which attention shapes behavior (Figure 2 and Methods for model details). We treated a Random Switching (RS) model as a baseline, which generates attentional trajectories without any learning or structured updating. This baseline allowed us to establish the minimum decoding performance expected in the absence of goal-directed attention. Although models from both classes successfully learned the task (Figure 2B), visual inspection of the latent representations suggest that, as expected, they rely on distinct assumptions about attention learning. FRL (Figure 2C)^4,8^ implements gradual value-driven updating attention, while SHT (Figure 2D)^5–7^, displays rapid, discrete shifts in attention through hypothesis switching. Together, these models instantiate qualitatively distinct assumptions about how attention is allocated during multidimensional RL. To determine which mechanism best captures human attention learning, we adopted a generative decoding framework that leverages the precise mechanisms specified by cognitive models and combines them with neural network approaches capable of decoding covert cognitive states.

**Figure 2.**
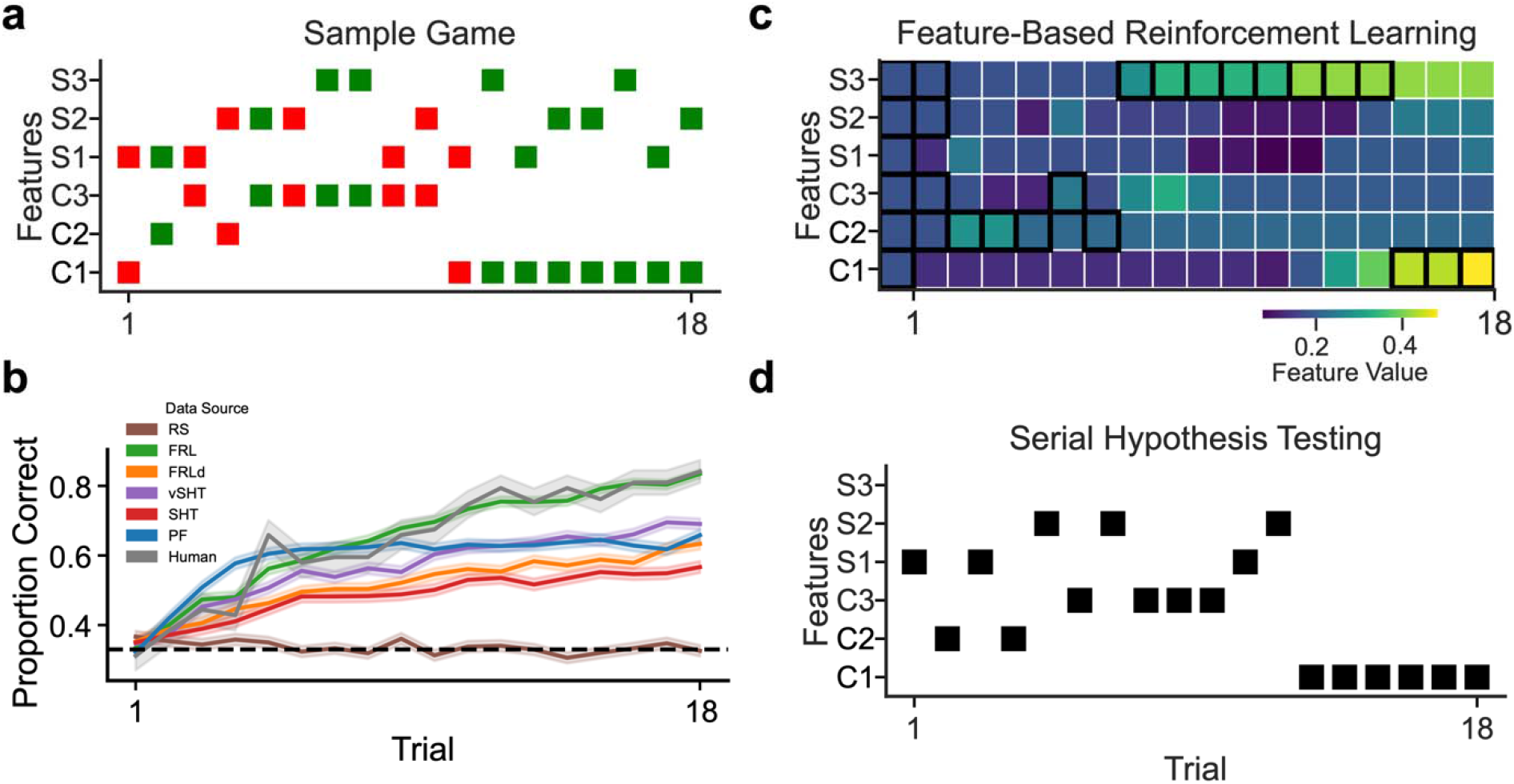
Cognitive models. **A**. Example choice game from the self-report human dataset (red = unrewarded, green = rewarded; features C1-3 correspond to colors and S1-3 to shapes). The correct target feature in this particular game was C1. Visual inspection suggests frequent attention switching early in the game, followed by convergence on the correct target feature (C1) in later trials. **B**. Proportion of correct choice increases across trials in simulated (N = 21; 40 games, 18 trial/game) and human (N = 21; 6 games, 18 trials/game; shading = SEM; dashed line = chance) data. For visual comparison to human data, 21 synthetic agents were selected from the 1000-agent simulated test dataset for each cognitive model. As expected, the Random Switching (RS) RS model shows no evidence of learning. However, all other models successfully learned the task, despite each proposing distinct attention learning dynamics, consistent with its underlying model class. Note that simulated agents were not fit to individual human behavior, parameters were just sampled from empirical priors, so the simulations are intended to illustrate qualitative learning dynamics rather than reproduce human performance quantitatively. **C**. Simulated feature values learned by the feature-based RL (FRL) model given the stimuli, choices, and rewards from the example game in panel A. The black square indicates the model’s inferred hypothesis on each trial (argmax over learned feature values). Consistent with gradual value updating, the FRL model switches its inferred hypothesis several trials after the participant’s apparent switch and maintains attention on a single feature for extended periods while updating values for all features on each trial. **D**. Trial-by-trial hypothesis inferred by the random serial hypothesis testing (rSHT) model for the same game. In contrast to FRL, rSHT maintains a single active hypothesis at a time and exhibits more rapid switching between features.

For each model, we trained a corresponding LaseNet Estimator^19^ to decode trial-by-trial feature-level attentional focus, consistent with the attentional dynamics assumed by that model (Figure 1C/D; see Methods). These models fall into two broad classes: those in which attention evolves gradually through learning, akin to RL processes, and those in which attention shifts rapidly via discrete hypothesis testing (Figure 1B).

The trial-by-trial attentional features decoded by each network provide an otherwise latent readout of the cognitive strategies that support adaptive learning in multidimensional environments.

### All networks recover latent attention above chance in synthetic data

After training six separate LaseNet Estimators^19^ for each of the cognitive models, we evaluated the decoding performance each network on two held-out test datasets. The first was a held-out test set simulated from the same model (N = 1000 agents per model; Figure 3). All networks performed well above chance at labeling the feature- and dimension-level attentional states generated from synthetic data from each respective cognitive model (feature-level attention mean accuracy (M ± SD): RS = 0.71 ± 0.17, FRL = 0.95 ± 0.02, FLRd = 0.94 ± 0.04, Hybrid = 0.88 ± 0.07, rSHT = 0.78 ± 0.15, PF = 0.86 ± 0.02, Figure 3A; dimension-level attention mean accuracy (M ± SD): RS = 0.72 ± 0.16, FRL = 0.96 ± 0.02, FLRd = 0.95 ± 0.03, Hybrid = 0.88 ± 0.07, rSHT = 0.78 ± 0.15, PF = 0.86 ± 0.02, Figure 3B). Analysis of single-game accuracy revealed that the networks often achieved perfect prediction in a varying proportion of games (FRL= 0.61; FLRd= 0.56; PF= 0.08; Hybrid= 0.27; rSHT= 0.21; RS= 0.14; 18 trials; Supplementary Figure 1), contributing to the high average decoding accuracy. Additionally, we observed no feature-specific biases in labeling accuracy across networks (Figure 3C). Taken together, these results validate our approach and demonstrate that neural estimators can directly learn to infer attention from behavior without requiring traditional model fitting to choice data.

**Figure 3:**
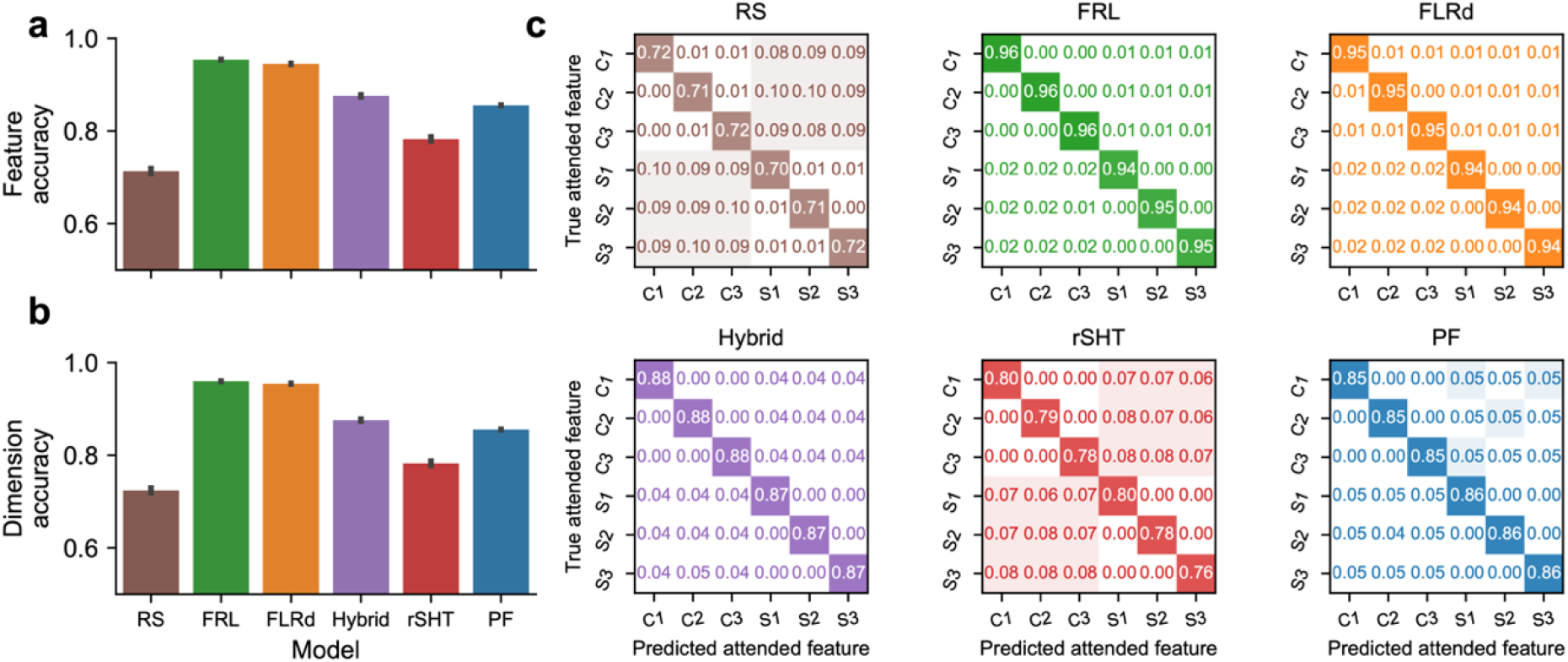
Evaluation of LaseNet Estimators on synthetic data. **A**. Attended feature labeling accuracy (chance = 0.167; error bars = SEM) for LaseNet Estimators evaluated on synthetic test data (N = 1,000 simulated agents/model, 40 games/agent, 18 trials/game). Feature accuracy reflects correct prediction of the attended feature (mean accuracy (M ± SD): RS = 0.71 ± 0.17, FRL = 0.95 ± 0.02, FLRd = 0.94 ± 0.04, Hybrid = 0.88 ± 0.07, rSHT = 0.78 ± 0.15, PF = 0.86 ± 0.02; error bars denote SEM). All networks performed well-above chance at attended feature decoding. **B**. Attended dimension labeling accuracy (chance = 0.50; error bars = SEM) for LaseNet Estimators evaluated on synthetic test data (N = 1000 agents; 40 games, 18 trials/game). Dimension accuracy reflects correct prediction of the attended dimension (shape vs. color), irrespective of specific feature identity (accuracy (M ± SD): RS = 0.72 ± 0.16, FRL = 0.96 ± 0.02, FLRd = 0.95 ± 0.03, Hybrid = 0.88 ± 0.07, rSHT = 0.78 ± 0.15, PF = 0.86 ± 0.02; error bars = SEM). All networks performed well-above chance at attended dimension decoding. **C**. Hypothesis labeling accuracy by feature for each cognitive model across simulated agents (N = 1000 simulated agents/model; chance = 0.167).

### Network decoding accuracy depends on cognitive model alignment

We first assessed how accurate each network was in predicting attention from behavioral trajectories generated by its own underlying cognitive model. Because each network was trained on synthetic data sampled from a specific generative model, we evaluated performance on held-out, unseen datasets generated by the same model to verify that decoding accuracy reflected successful recovery of the intended latent cognitive structure. This within-class generalization test allowed us to verify that each network had internalized the characteristic attentional dynamics of its training distribution before comparing performance across model classes or on human data (see Methods).

As expected, each model’s synthetic data were labeled most accurately by the network trained on that same model (see Methods; Supplementary Figure 3), indicating that these networks are not general-purpose decoders but instead preferentially capture attentional strategies specific to their generative assumptions.

We next investigated the generalizability of the FRLd and Hybrid networks for within- and between-class cognitive models. Specifically, we tested whether estimators trained under the FRLd and Hybrid models could accurately label hypotheses in synthetic datasets generated by all candidate models (see Methods; Figure 4), or whether they instead preferentially captured model-specific feature–attention switching dynamics. Both the FRLd and Hybrid networks achieved their highest decoding accuracy when evaluated on data generated by the same cognitive model on which they were trained, with reduced performance on data generated by alternative model classes (FRLd network accuracy (M ± SD): RS evaluation data = 0.37 ± 0.22, FRL evaluation data = 0.67 ± 0.10, FRLd evaluation data = 0.94 ± 0.04, Hybrid evaluation data = 0.76 ± 0.09, rSHT evaluation data = 0.62 ± 0.19, PF evaluation data = 0.61 ± 0.04; Figure 4A; Hybrid network accuracy (M ± SD): RS evaluation data = 0.47 ± 0.27, FRL evaluation data = 0.57 ± 0.14, FRLd evaluation data = 0.78 ± 0.09, Hybrid evaluation data = 0.88 ± 0.07, rSHT evaluation data = 0.76 ± 0.16, PF evaluation data = 0.76 ± 0.03, Figure 4B).

**Figure 4.**
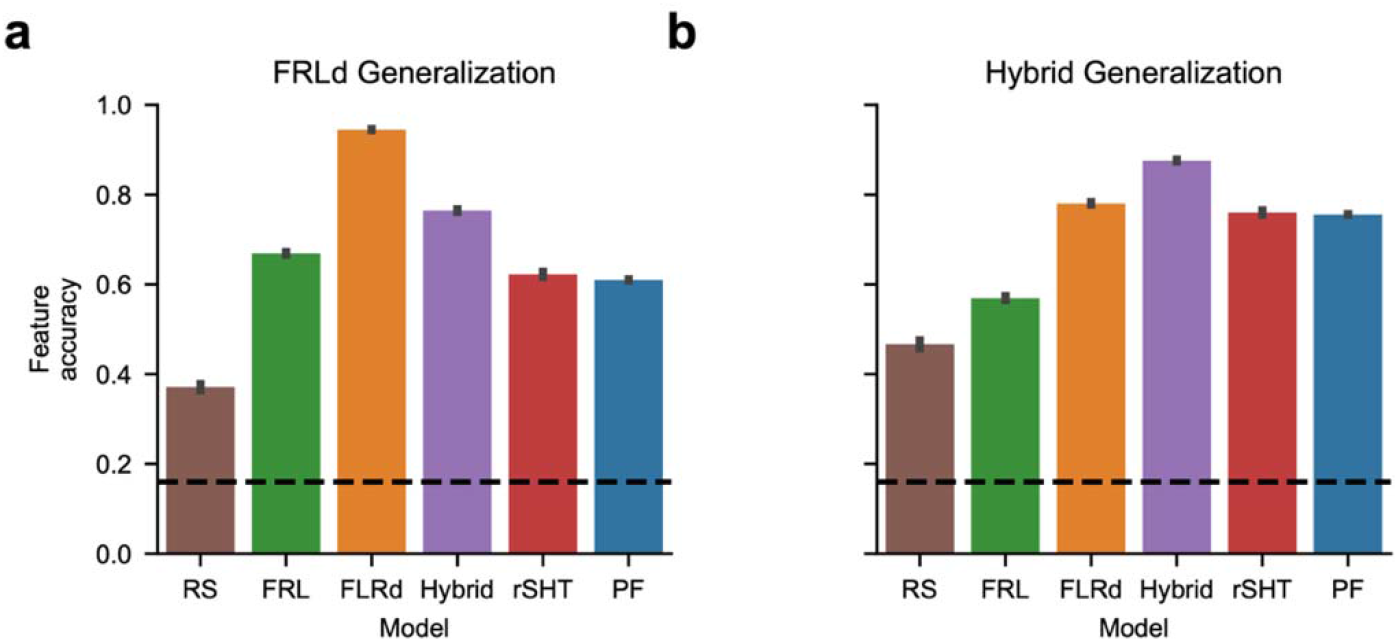
Network generalization across generative models. **A**. The FRLd-trained network generalized best to synthetic test data generated from the FRLd generative model. It showed the next-highest performance on data generated from the Hybrid model, of which FRLd is a constituent, and third-highest performance on data generated from the FRL model, which belongs to the same feature-based RL model class (FRLd network accuracy (M ± SD): RS evaluation data = 0.37 ± 0.22, FRL evaluation data = 0.67 ± 0.10, FRLd evaluation data = 0.94 ± 0.04, Hybrid evaluation data = 0.76 ± 0.09, rSHT evaluation data = 0.62 ± 0.19, PF = 0.61 ± 0.04; N = 1,000 simulated agents/model, 40 games/agent, 18 trials/game; error bars =SEM). **B**. The Hybrid-trained network generalized best to synthetic test data generated from the Hybrid generative model. It generalized second best to data generated from the FRLd model, one of its constituent components, and showed similar third- and fourth-best performance on data generated from SHT-class models (Hybrid network accuracy (M ± SD): RS evaluation data = 0.47 ± 0.27, FRL evaluation data = 0.57 ± 0.14, FRLd evaluation data = 0.78 ± 0.09, Hybrid evaluation data = 0.88 ± 0.07, rSHT evaluation data = 0.76 ± 0.16, PF= 0.76 ± 0.03; N = 1,000 simulated agents/model, 40 games/agent, 18 trials/game; error bars =SEM).

This pattern indicates that networks trained under distinct cognitive assumptions learn model-specific representations of attentional dynamics, rather than a single, model-agnostic decoding strategy, and that these representations do not fully generalize across model types. This highlights the interpretability of the assumptions embedded in each network when inferring latent cognitive variables.

### Human attention learning is best described by an interaction between value updating and hypothesis testing

Having shown that the networks can adequately decode synthetic data, we next evaluated each network’s ability to label trial-by-trial attended features in a second held-out test dataset generated from human participants who reported their attentional focus on each trial but for whom their ground truth generative cognitive model is unknown (Figure 5).

**Figure 5:**
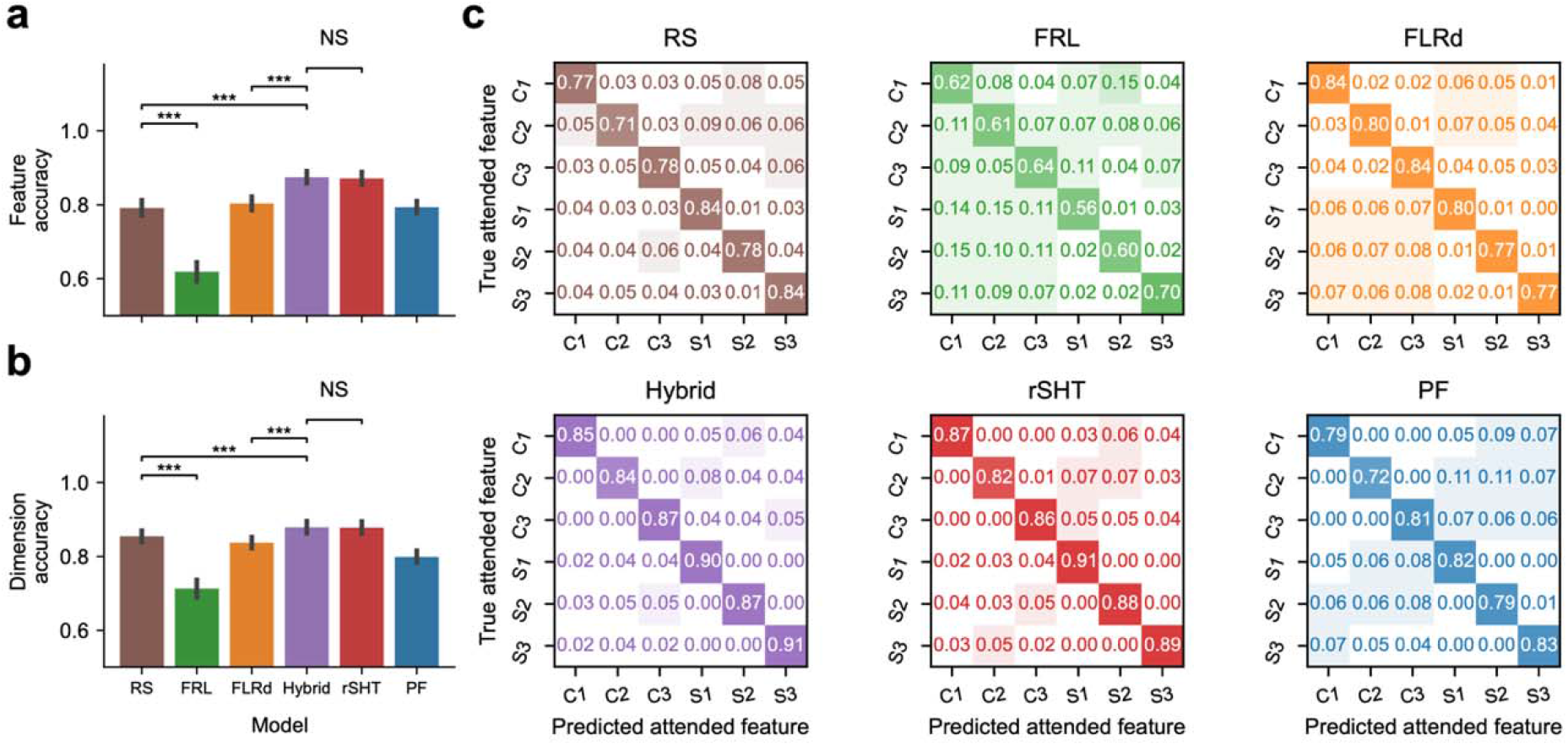
Evaluation of LaseNet Estimators on human self-report data. **A**. Attended feature decoding accuracy across LaseNet networks. Feature accuracy reflects correct prediction of the attended feature (mean accuracy (M ± SD): RS = 0.79 ± 0.09; FRL = 0.62 ± 0.12, FLRd = 0.80 ± 0.08, Hybrid = 0.87 ± 0.08, rSHT = 0.87 ± 0.08; PF = 0.79 ± 0.07; chance accuracy = 0.167; error bars = SEM). All networks performed well-above chance at attended feature decoding. Shown are selected significant pairwise comparisons relative to the RS baseline and the Hybrid network (hypothesized to perform best); see Results for full statistical reporting. RS performed significantly worse than the Hybrid network (β = 0.08, *z* = 8.85, *p* < 0.001) but significantly better than the FRL network (β = −0.17, *z* = −8.99, *p* < 0.001). Among the Hybrid model and its constituent components, the Hybrid outperformed FRLd (FRLd – Hybrid: Δ = -0.07, *z* =- 6.40, *p* < 0.001; FDR corrected), but did not differ significantly from rSHT in attended feature decoding (Hybrid – rSHT: Δ = 0.003, *z* = 0.31, *p* = 0.75; FDR corrected). See Results for full pairwise comparison results. **B**. Dimension accuracy reflects correct prediction of the attended dimension (shape vs. color), irrespective of specific feature identity (mean accuracy (M ± SD): RS = 0.85 ± 0.07; FRL = 0.71 ± 0.11, FLRd = 0.84 ± 0.07, Hybrid = 0.88 ± 0.08, rSHT = 0.88 ± 0.08, PF = 0.80 ± 0.07; chance accuracy = 0.50; error bars = SEM). All networks performed well-above chance at attended dimension decoding. Shown are selected significant pairwise comparisons relative to the RS baseline and the Hybrid network (hypothesized to perform best); see Results for full statistical reporting. RS performed significantly worse than the Hybrid network (β = 0.03, *z* = 3.54, *p* < 0.001) but significantly better than the FRL network (β = −0.14, *z* = −5.35, *p* < 0.001). Among the Hybrid model and its constituent components, the Hybrid outperformed FRLd (FRLd – Hybrid: Δ = -0.04, *z* = -4.28, *p* < 0.001; FDR corrected), but did not differ significantly from rSHT in attended dimension decoding (Hybrid – rSHT: Δ = 0.001, *z* = 0.16, *p* = 0.87; FDR corrected). Together, these results indicate that incorporating SHT substantially improves decoding accuracy beyond gradual FRL alone. See Results for full pairwise comparison results. **C**. Hypothesis decoding accuracy by feature for each network across participants (N = 21; chance = 0.167).

These self-reports provided a ground-truth measure of otherwise covert attention for model evaluation. We first examined feature-level decoding accuracy, which was defined as whether the network’s predicted feature (i.e., “square”, or “blue) was the same as the true attended feature from the self-report human dataset in which the generative cognitive model is unknown (N = 21 participants; feature-level attention mean accuracy (M ± SD): RS network = 0.79 ± 0.09; FRL network = 0.62 ± 0.12, FLRd network = 0.80 ± 0.08, Hybrid network = 0.87 ± 0.08; rSHT network = 0.87 ± 0.08; PF network = 0.79 ± 0.07; Figure 5A).

Using a mixed-effects model (see Methods; Eq. 1), we tested whether each network decoded human self-reported attention more accurately than the RS baseline, which assumes no attention-learning mechanism. Relative to RS, the FRL network showed substantially worse decoding accuracy (β = −0.17, *z* = −8.99, *p* < 0.001; Figure 5A). The FRLd network did not exhibit significantly improved feature accuracy from the RS network (β = 0.01, *z* = 1.26, *p* = 0.21; Figure 5A). In contrast, several models significantly outperformed RS. The Hybrid (β = 0.08, *z* = 8.85, *p* < 0.001; Figure 5A) and rSHT (β = 0.08, *z* = 8.55, *p* < 0.001; Figure 5A) networks showed significant improvements in feature decoding accuracy. Notably, whereas FRLd did not yield a significant improvement over RS, augmenting it with SHT in the Hybrid model produced a substantially larger gain, isolating the contribution of value-based switching to decoding human attentional dynamics. The PF did not improve relative to RS (β = 0.002, *z* = 0.17, *p* = 0.86; Figure 5A). Together, these results indicate that models incorporating structured hypothesis switching substantially outperform RS in decoding attended features, whereas feature-based RL without decay performs significantly worse.

Next, we ran pairwise comparisons to compare performance among models that incorporated distinct attention-learning mechanisms (FRL, FRLd, rSHT, Hybrid, PF; see Methods). Pairwise contrasts revealed strong and systematic differences in feature decoding accuracy across models. Hybrid performed significantly better than all other networks besides rSHT (FRL – Hybrid: Δ =-0.26, *z* =-12.48, *p* < 0.001; FRLd – Hybrid: Δ = -0.07, *z* =-6.40, *p* < 0.001; Hybrid – rSHT: Δ = 0.003, *z* = 0.31, *p* = 0.75; Hybrid − PF: Δ = 0.08, *z* = 6.84, *p* < 0.001; FDR corrected; Figure 5A). The rSHT network performed significantly better than the networks in the FRL class as well as the PF network (FRL – rSHT: Δ = -0.25, *z* =-12.50, *p* < 0.001; FRLd – rSHT: Δ = -0.07, *z* = -6.12, *p* < 0.001; PF – rSHT: Δ = -0.08, *z* = -7.40, *p* < 0.001; FDR corrected; Figure 5A). The FRLd network performed significantly better than the FRL network (FRL – FRLd: Δ = -0.19, *z* = -10.60, *p* < 0.001; FDR corrected; Figure 5A). The PF network performed significantly better than the FRL network but did not perform differently than the FRLd network (FRL – PF: Δ = -0.18, *z* = -8.67, *p* < 0.001; FRLd – PF: Δ = 0.01, *z* = 0.71, *p* = 0.53; FDR corrected;

Figure 5A). Overall, these comparisons reveal a consistent performance ordering in which Hybrid and rSHT networks performed best, followed by PF and FRLd, with the basic FRL network performing worst.

We next examined decoding accuracy against the RS baseline at the attended dimension level using an analogous mixed-effects model (see Methods, Eq. 1). Dimension-level decoding accuracy was defined as whether the network’s predicted feature belonged to the same dimension (shape or color) as the true attended feature from the self-report human dataset, quantifying accurate inference of the attended dimension independent of specific feature identity (N = 21 participants; dimension-level attention mean accuracy (M ± SD): RS network = 0.85 ± 0.07; FRL network = 0.71 ± 0.11, FLRd network = 0.84 ± 0.07, Hybrid network = 0.88 ± 0.08, rSHT network = 0.88 ± 0.08, PF network = 0.80 ± 0.07, Figure 5B). Relative to RS, FRL again performed significantly worse (β = −0.14, *z* = −5.35, *p* < 0.001; Figure 5B). The FRLd model did not differ from RS (β =-0.02, *z* = −1.72, *p* = 0.09; Figure 5B), indicating comparable dimension-level decoding performance. In contrast, both Hybrid (β = 0.03, *z* = 3.54, *p* < 0.001; Figure 5B) and rSHT (β = 0.02, *z* = 2.63, *p* = 0.009; Figure 5B) showed significant improvements over RS. Together, these results suggest that incorporating SHT more effectively captures the structured, dimension-level organization of attention learning in human behavior than FRL alone. The PF model performed significantly worse than RS at the dimension level (β = −0.06, *z* = −3.77, *p* < 0.001; Figure 5B). Overall, these results mirror the feature-level findings, with rSHT and Hybrid models outperforming RS, highlighting the utility of sequential hypothesis testing for capturing the fast-attention switching dynamics characteristic of human behavior, and FRL showing consistent deficits in attention decoding.

Pairwise contrasts for dimension decoding accuracy (see Methods) revealed a similar pattern. Hybrid performed significantly better than all other networks besides rSHT (FRL – Hybrid: Δ = -0.17, *z* = -6.25, *p* < 0.001; FRLd – Hybrid: Δ = -0.04, *z* = -4.28, *p* < 0.001; Hybrid – rSHT: Δ = 0.001, *z* = 0.16, *p* = 0.87; Hybrid − PF: Δ = 0.08, *z* = 6.74, *p* < 0.001; FDR corrected; Figure 5B). The rSHT network performed significantly better than the networks in the FRL class as well as the PF network (FRL – rSHT: Δ = -0.17, *z* = -6.21, *p* < 0.001; FRLd – rSHT: Δ = -0.04, *z* = -3.68, *p* < 0.001; PF – rSHT: Δ = -0.08, *z* = -7.82, *p* < 0.001; FDR corrected; Figure 5B). The FRLd network performed significantly better than the FRL network (FRL – FRLd: Δ = -0.12, *z* = -5.46, *p* < 0.001; FDR corrected; Figure 5B). The PF network performed significantly better than the FRL network but significantly worse than the FRLd network (FRL – PF: Δ = -0.09, *z* = -3.53, *p* < 0.001; FRLd – PF: Δ = 0.04, *z* = 2.50, *p* < 0.05; FDR corrected; Figure 5B). Additionally, no dimension-specific biases in labeling accuracy were observed across networks for the human self-report data (Figure 5C). Together, these results indicate that incorporating SHT substantially improves decoding performance compared to FRL alone.

Importantly, Hybrid decoding accuracy was relatively stable across participants, remaining high for both the best- and worst-performing individuals (Figure 6), and exceeding chance levels even in cases with lower labeling accuracy (Figure 6). In addition, the Hybrid model qualitatively reproduced human-like patterns of attention switching (Supplementary Figure 2).

**Figure 6:**
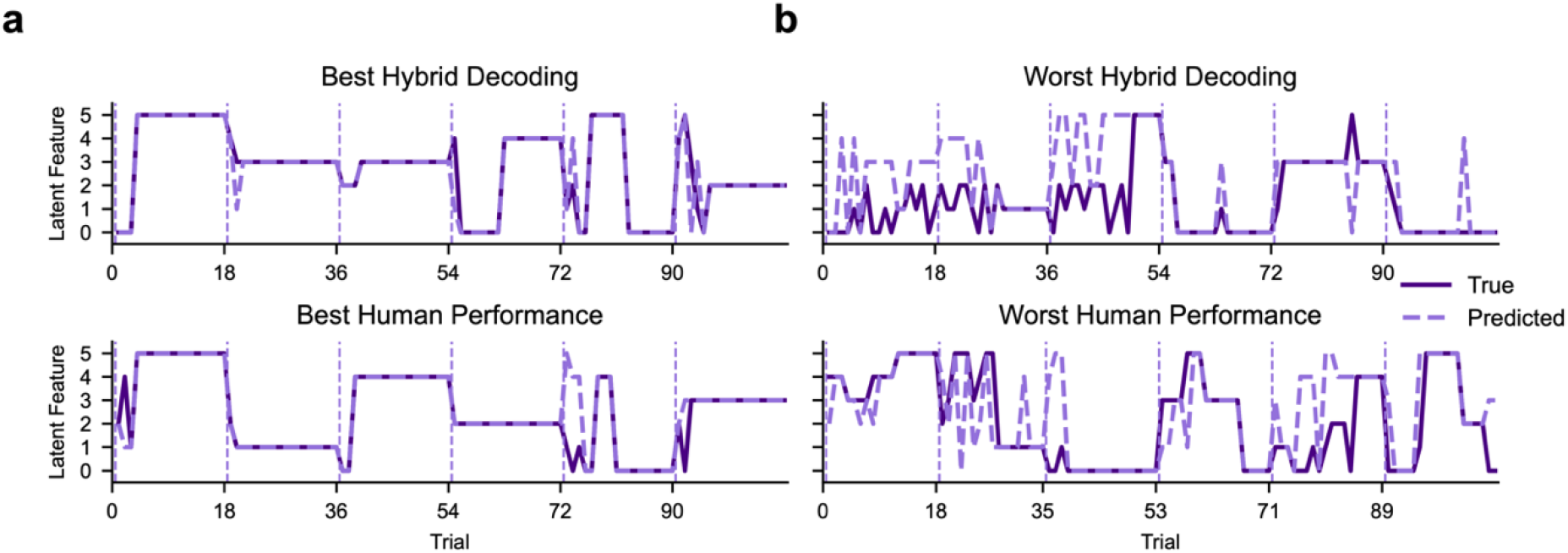
Representative examples of Hybrid network predictions for the best- and worst-decoded participants. **A**. The highest Hybrid decoding accuracy (top; 0.72 behavioral accuracy; 0.95 Hybrid Network decoding accuracy) alongside the best-performing participant by behavioral accuracy (bottom; 0.92 behavioral accuracy; 0.95 Hybrid Network decoding accuracy). **B**. The lowest Hybrid decoding accuracy (top; 0.60 behavioral accuracy; 0.69 Hybrid Network decoding accuracy) alongside the worst-performing participant (bottom; 0.36 behavioral accuracy; 0.73 Hybrid Network decoding accuracy). Note, the lowest performing participant missed one trial. This trial was excluded from the visualization, resulting in 107 trials shown for this participant rather than the standard 108 trials displayed for the other participants. The dark purple solid trace indicates the attended feature reported by participants, while the light purple dashed trace shows the Hybrid network’s predictions. Notably, the Hybrid Network did not perform worst when labeling the behaviorally poorest-performing participant and achieved comparable decoding accuracy across participants with different levels of task performance, indicating that latent-state decoding accuracy was not tightly coupled to overt behavioral performance.

While the rSHT and Hybrid models both exhibited superior attended-feature and - dimension decoding accuracy compared to the other networks, they did not outperform each other (Figure 5A/B). To investigate whether differences between these models emerge at a more granular level of inference, we examined whether the Hybrid network captures additional structure in the distribution of candidate hypotheses underlying attentional switches (Figure 7).

**Figure 7:**
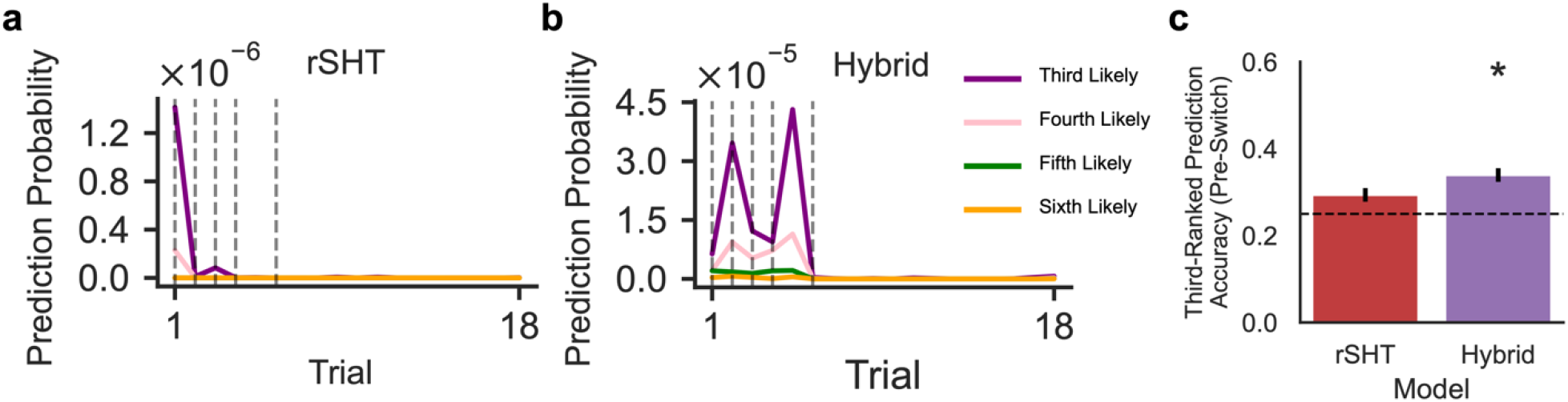
Hybrid network learns a human-like distribution over alternative hypotheses preceding switches. Example game illustrating the trial-resolved likelihood assigned to non-selected features by the **A**. rSHT and **B**. Hybrid networks. Traces show the predicted probability assigned to the 3rd–6th most likely features on each trial. Dashed vertical lines denote ground-truth attentional switches based on participant’s self-report. The Hybrid network maintains a broader distribution over candidate features, consistent with informed hypothesis switching. **C**. Group-level analysis of hypothesis structure preceding attentional switches. For each participant, switch trials were isolated, and accuracy was computed for the third-ranked predicted feature on the trial immediately preceding a switch to a new shape or color (i.e., not selected on the preceding trial). This rank was selected because the top-ranked features correspond to the currently attended stimulus dimensions, enabling assessment of model representations of alternative candidate hypotheses. Only the Hybrid network showed significantly above-chance accuracy in identifying the subsequently attended feature (one-sample t-test: t = 2.65, p < 0.05). Bars indicate group means, error bars denote SEM, and the dashed line marks chance performance.

To do this, we analyzed the trial-by-trial probability assigned by each network to alternative candidate features, focusing on trials immediately preceding attentional switches. We hypothesized that networks trained on data more closely aligned with human switching dynamics would exhibit belief distributions that more strongly weight the feature participants subsequently switched to. In other words, prior to a switch, such models should already assign elevated probability to the “switched-to” hypothesis.

In an example game, the rSHT network sharply collapsed probability mass onto the currently selected feature, assigning near-zero probability to alternatives (Figure 7A). In contrast, the Hybrid network which maintained a broader distribution over multiple plausible hypothesis (Figure 7B), consistent with its assumption that switching is informed by RL values that decay gradually for unselected features.

To quantify this effect at the group level, we isolated switch trials for each participant and examined the model’s probability ranking of the feature that was subsequently selected, focusing on the trials immediately preceding a switch to a feature not chosen on the prior trial. Specifically, we evaluated whether this feature was ranked third in the model’s probability distribution. The third rank was selected because the top-ranked features correspond to the current choice, allowing us to assess the representation of alternative hypotheses beyond the chosen features.

Across participants, only the Hybrid network ranked the subsequently selected feature above chance levels (one-sample t-test: *t* = 2.65, *p* < 0.05; Figure 7C). This finding supports the Hybrid model’s prediction that attentional switches are informed by a learned distribution of feature values accumulated through experience.

Notably, this effect emerged despite comparable overall decoding accuracy across models. The Hybrid network uniquely captured graded, anticipatory belief updating over alternative hypotheses, more closely mirroring how humans weight candidate task states prior to shifting attention. Thus, it is the internal structure of the Hybrid model’s belief representation – not merely its final output – that aligns with human attentional dynamics. Collectively, these findings indicate that the Hybrid network best captures key features of human switching behavior and generalizes across individuals with heterogeneous task performance.

## Discussion

A central challenge in human learning and decision-making is how selective attention enables RL to remain tractable in high-dimensional, partially observable environments ^2,16,21,23^. Natural settings rarely provide an explicit mapping between sensory input and task-relevant state. Instead, individuals must determine which features warrant attention and which can be ignored. Although RL is widely regarded as a normative framework for adaptive behavior, it presupposes a compact and task-relevant state representation. Without a mechanism for filtering irrelevant dimensions, the learning problem quickly becomes intractable and fails to capture real-world cognition. Therefore, understanding how attention filters and structures environmental input is essential for establishing RL as a mechanistic account of human behavior.^2,16,21^.

In this work, we addressed a fundamental gap in the study of attention learning: although competing computational models posit distinct mechanisms for attention learning, covert attentional states are not directly observable and are therefore non-identifiable from choice data alone. By combining mechanistic cognitive models with LaseNet^19^, a novel RNN-based approach capable of recovering latent cognitive dynamics from behavior, we moved beyond indirect inference through choice fitting. Instead, we directly decoded trial-by-trial attentional states under competing cognitive strategies, enabling adjudication between alternative theories of attention learning.

We trained RNNs using LaseNet on synthetic datasets generated from six distinct cognitive models (FRL, FRLd, rSHT, Hybrid, PF, RS) to infer trial-by-trial attentional allocation. Rather than comparing these cognitive models based on their ability to reproduce observed choices, we use each as a source of structured inductive biases during training, allowing separate RNNs to encode different assumptions about how attention is learned. This approach enables direct comparison of how well each model can infer attention across synthetic and human test data, circumventing difficulties caused by traditional cognitive model fitting methods based on MLE^24^.

We found that all networks performed well above chance in inferring attention on simulated test data. To rule out the possibility that the Hybrid model serves as a general-purpose decoder, we tested its ability to label latent variables in datasets generated by other cognitive models. Interestingly, as hypothesized, the Hybrid network did not generalize well to out-of-class synthetic data, performing best only on data generated by its own generative process. Further, the Hybrid network best captured human attention learning dynamics. This result, together with the Hybrid network’s limited generalizability to data generated by other cognitive models, suggests that this model captured a distinct algorithmic bias that most closely aligns to human attention dynamics.

Interestingly, while FRL-trained networks predict their own attended feature switching behavior with high accuracy in synthetic data, they perform worse than SHT-alone and Hybrid networks when applied to infer latent attentional state in human data. This pattern suggests that, while FRL networks fit their own data well, they do not encode attentional dynamics that align with human cognitive strategies. This likely reflects a strong inductive match between the FRL training and synthetic test distributions, but weaker alignment between FRL-generated behavior and the attentional dynamics observed in human data. Our work demonstrates that accurately decoding human attention dynamics requires not just a high-performing network but also training on synthetic data from cognitive models with the appropriate mechanistic constraints.

In contrast, the Hybrid network, trained on data that integrates both FRL and SHT mechanisms exhibits strong generalization to human data. This suggests that the Hybrid dataset imparts inductive biases that are robust across behavioral regimes and share structure with human cognition. The superior alignment of the Hybrid network with human attentional dynamics suggests that human state representation learning reflects an integration of gradual value-based updating and rapid hypothesis switching. Notably, the Hybrid model had previously been identified as the best account of human choice behavior in a similar task environment^6^. Here, we show that the same algorithmic framework also provides the most accurate account of latent attentional allocation. This convergence across observable choice and inferred attention suggests that the Hybrid model captures a unified computational strategy underlying both decision outcomes and the internal dynamics of representational processes that support them.

One strength of our approach was the use of informed free parameter priors for generating both training and test data, derived from previous studies. These priors helped to constrain the parameter space more effectively, improving the network’s ability to capture realistic patterns in the data. One important design choice, and potential limitation, was the absence of model-specific hyperparameter tuning prior to training. All networks shared the same architecture, input–output structure, and training schedule, and were trained on datasets with identical statistical structure. The sole systematic difference across training sets was the attentional trajectories generated by the underlying representation learning models. This design ensured that any differences in network performance could be attributed to the inductive biases embedded in the training data, rather than to differences in model capacity, optimization, or architectural choices.

While this approach means that individual networks may not be fully optimized for their respective synthetic datasets, the primary goal of this work was not to maximize performance for any single model class, but to assess how different cognitive strategies, when used to generate training data, influence a network’s ability to extract latent cognitive variables from behavior. Since the structure and dimensionality of the training data were held constant across models, we do not expect substantial qualitative differences to emerge from extensive hyperparameter tuning. Nonetheless, future work could explore whether model-specific tuning yields incremental gains in latent variable inference without altering the core conclusions.

A notable finding was the performance of the RS network in labeling human data, particularly its ability to outperform the FRL model. Although the RS network does not explicitly model learning dynamics underlying adaptive task performance, it captures rapid attentional switching behavior that more closely resembles human behavior than the gradual updating characteristic of FRL. This result highlights the importance of fast attentional dynamics for state representation learning and underscores the limitations of RL alone in accounting for the rapid shifts in attention that support learning in multidimensional tasks.

More broadly, our findings highlight how theory-informed generative cognitive models can do more than predict choices: they can serve as structured training signals for general-purpose neural networks, enabling recovery of otherwise unobservable cognitive dynamics. By using cognitive theory to generate training data, we show that theoretical commitments can directly shape the internal representations learned by neural decoders. This theory-informed RNN framework demonstrates how combining generative cognitive models with flexible function approximators moves beyond traditional maximum-likelihood model comparison linking computational algorithms to covert, trial-by-trial attentional dynamics. Such approaches offer a principled path for inferring latent mental states and adjudicating competing theories of learning in complex, high-dimensional environments.

## Methods

### Multidimensional RL Task

Our training environment for attention decoding was a multidimensional RL task adapted from prior work^4,5,8–10^. In the human experiment, participants (N = 21; age M ± SD = 32 ± 14.65) completed six 18-trial games in which they made repeated choices between three stimuli varying in shape (square, oval, circle) and color (orange, yellow, blue; Figure 1A). Each game had one relevant dimension (shape or color) and a target feature (e.g., square). After each choice, participants were rewarded with 80% probability if they selected the stimulus containing the target feature, and with 20% probability otherwise. To maximize reward, participants had to learn the target feature and relevant dimension via trial-and-error. Changes in the relevant dimension were explicitly signaled between games. And participants were aware of the generative structure of the task (i.e. one target feature, one dimension being relevant, and the exact reward probabilities).

Because human participants’ “ground truth” attention is latent in this task^4–6,8^, we collected a labeled dataset of self-report human test data (N = 21 participants; 6 games/participant, 18 trials/game, 108 trials in total) in which the participant was instructed to self-label their focus of attention on each trial. Participants were asked to record audio of their belief about the target feature on each trial. These audio recordings were then transcribed and used for network evaluation, referred to throughout as the “self-report” human dataset. We reasoned that because participants are aware of the generative structure of the task (i.e. there is one target feature within a relevant dimension), they should be able to verbalize where they are focusing their attention on each trial. Although we have access to this self-report information, we do not have access to the generative model that produced their choices. Critically, this human test dataset allows us to assess the ability of LaseNet Estimators^19^ trained on synthetic data from different cognitive models to infer the participants’ latent mechanism of attention allocation.

### Cognitive Models

Two cognitive model classes have been proposed as mechanisms of attention allocation during representation learning: FRL^4,11–15^ and SHT^6–8,17^. Models within these classes differ in their assumed attentional dynamics: FRL models update attention gradually through learning, whereas SHT models implement rapid, discrete switching (Figure 2). A Hybrid model that combines elements of both FRL and SHT, has also been proposed and was previously shown to best capture participants’ choice behavior during multidimensional RL^6^. Models within these respective classes seek to explain how attention learning mechanisms support decision-making by prioritizing relevant information during learning.

### Feature Reinforcement Learning (FRL)

FRL models (Algorithm 1) maintain values (*v*) for each of the six features (denoted by *f*_*i,j*_ where *i* represents the dimension and *j* represents the feature). The baseline FRL model assumes that for each trial *t*, the expected value (*EV*) of each stimulus (*s*) is calculated as the sum of its feature values. The choice probability is computed using a noisy softmax, with inverse temperature (β) as a free parameter. On each trial, the action *c*_*t*_ is sampled from this probability distribution. Feature values *v*_*t*_ (*f*_*i,c*_) for the chosen stimulus (*c*_*t*_) are updated according to a Rescorla-Wagner update rule^25^, with learning rate (η) as a free parameter. The FRL with decay model^8^ maintains the same learning rule as the FRL model for updating feature values associated with the chosen stimulus (*c*_*t*_), with an additional decay term to update values for unchosen features. Under this model, feature values for unchosen stimuli decay towards zero with a factor of (where is between 0 and 1). β, η, and are free parameters. For both models, the hypothesis on any given trial corresponds to the feature with the highest value following value update.

#### Algorithm 1

Feature-Based Reinforcement Learning (FRL) Model Class

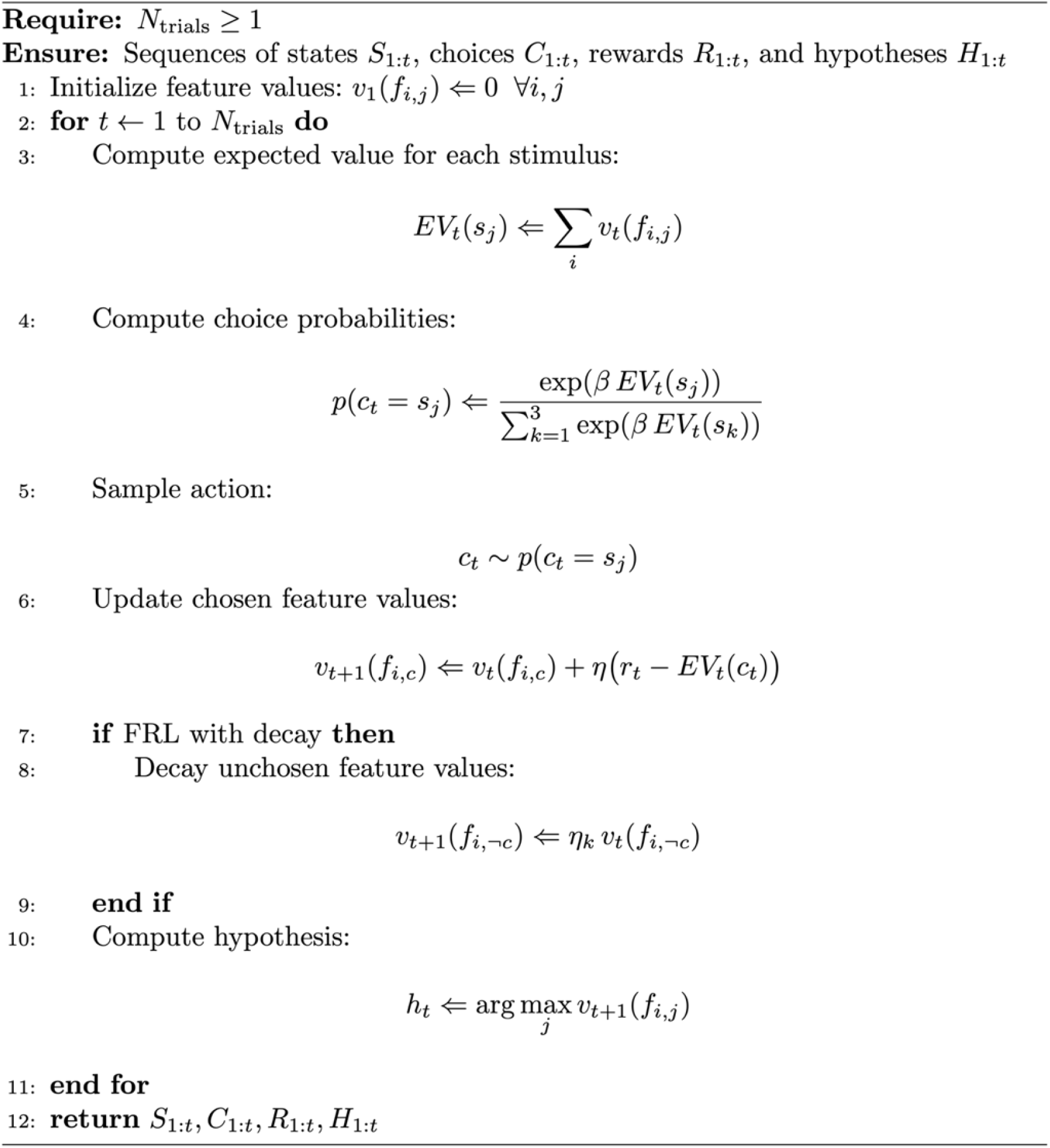

### Serial Hypothesis Testing (SHT)

SHT models (Algorithm 2) assume that participants attend to one hypothesis () regarding the target feature on any given trial. The generative model contains a hypothesis-testing policy (whether to stay with the current hypothesis or switch to a new one) and a hypothesis-switching policy (what the next hypothesis should be when switching hypotheses). Different variants of this model have been proposed that primarily differ in the hypothesis switching policy. The random SHT model (rSHT)^5^ assumes that on each trial the participant estimates the reward probability of the current hypothesis (ρ_*reward*_). The estimated reward probability is compared to a soft threshold (θ), to determine whether to stay with the current hypothesis or randomly switch to one of the other possible hypotheses. θ and β_stay_ are free parameters.

The particle filter model (PF)^7^ assumes beliefs about the target feature are updated using memory-augmented particle filtering. Like the rSHT model, the PF maintains a single hypothesis about the target feature. To decide how to switch, the PF considers the current choice and reward, and the previous *n* choices and rewards stored in memory. Based on these observations, it computes a proposal distribution representing the probability of staying with the current hypothesis or switching to one of the other features. The proposal distribution represents the model’s belief about the target feature, given the hypothesis and evidence on the previous *n* trials. Thus, the PF accounts for limited working memory constraints, but uses memory to compute an optimal switch probability given its beliefs. Here, the agent’s hazard rate (the probability that the target feature can change on a given trial) and *n* (the amount of evidence retained in working-memory), are free parameters.

Finally, we considered a Hybrid model that combines elements of SHT with FRL^6^. Like the rSHT model and the PF, the Hybrid model maintains a single hypothesis about the target feature. To decide how to switch, the Hybrid model uses a value-based hypothesis-switching policy updated based on FRL with decay (Algorithm 1). Here, β_*switch*_ is an inverse temperature parameter that modulates the sensitivity of attention switching toward the feature with the highest learned value according to FRLd. The Hybrid model can be thought of approximating the proposal distribution using FRL with decay (rather than optimal inference, as in the case of the PF model).

#### Algorithm 2

Serial Hypothesis Testing (SHT) Model Class

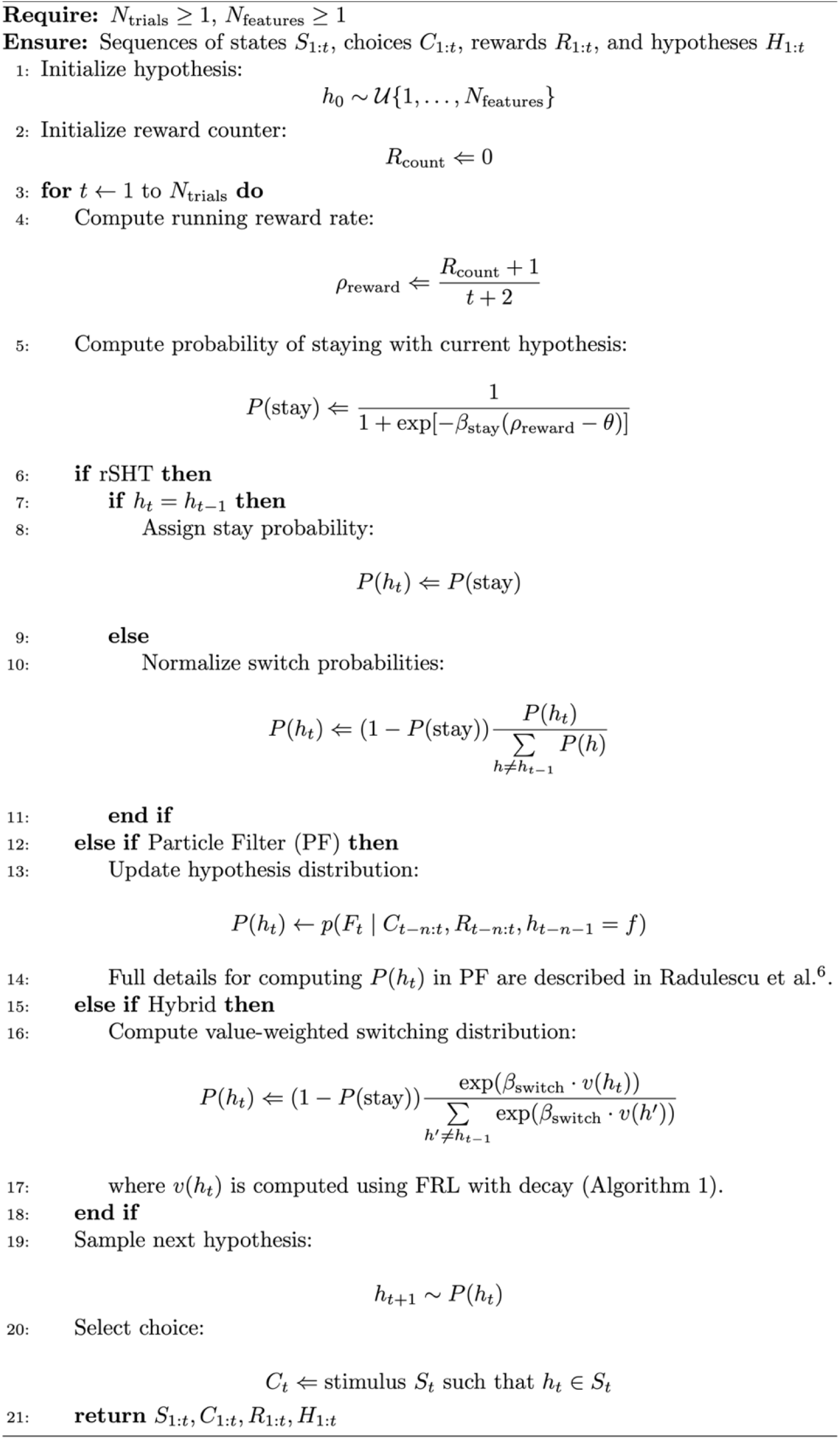

### Random Switching (RS)

As a final comparison, we trained a model which stochastically switched attention between features with a fixed lapse rate, denoted by λ, on each trial. Unlike the other models, the RS model did not update feature values based on rewards or maintain any structured representation of the task. Instead, on each trial, the attended feature was either maintained with probability 1 − λ or randomly switched to a new feature with probability λ, where λ was drawn from a uniform distribution between 0 and 1. The RS model served as a baseline, allowing us to evaluate how a LaseNet Estimator^19^ trained on a cognitive model that lacks systematic learning and hypothesis testing based on evidence would perform.

### LaseNet Estimators

Trial-by-trial attention decoding in the multidimensional RL task was done on synthetic data generated from six cognitive models. Throughout, we refer to these trained models using the convention *cognitive model name + network* (e.g., Hybrid network), indicating the LaseNet instance trained under that specific cognitive model. To decode attention, we used LaseNet^19^, a novel RNN-based method which directly maps choice data to latent variable space. Unlike traditional MLE methods, which require fitting model parameters for model comparison and inference^26^, LaseNet directly infers latent variables from behavior, and is thus well suited for models with analytically intractable or computationally intensive likelihoods. The LaseNet architecture consists of a bidirectional gated recurrent network that captures both past and future context around each decision point, followed by multilayer perceptrons that map context-aware embeddings to continuous and/or discrete latent variable spaces. The result is a flexible framework for decoding latent cognitive dynamics from high-dimensional behavioral sequences, enabling high-precision comparisons between cognitive models not just in terms of choice predictions, but in terms of their ability to explain how latent cognitive variables evolve over time.

### Training

Our goal was to infer participants’ attention allocation by decoding the feature (i.e., “yellow”) they were attending to on each trial. During the training phase (Figure 1C, adapted from Pan et al.^19^, we created a synthetic dataset by simulating the desired cognitive model (see Cognitive Models section). Each pair was generated with informed priors θ, by sampling parameter values uniformly between the minimum and maximum of the fitted ranges reported in prior studies^5–8^. LaseNet was trained using model-simulated observable data as input and a series of model-derived latent variables (i.e. attended feature) as output. For each LaseNet estimator, we simulated 20,000 (*Z, Y*) pairs, with 720 trials in each pair, as training data. In effect, this means that each individual parameter setting (“agent”) generated 40 games, each representing an instance of dynamic attention allocation under a specific cognitive model. Notably, while FRL models produce fixed attention trajectories under the same parameter setting, SHT models update attention stochastically, meaning each game represents a possible trajectory through hypothesis space (see Cognitive Models section). Because our goal was to compare how the algorithmic biases imparted by each cognitive model influence decoding accuracy, we intentionally used a fixed architecture and training protocol across all networks to isolate the influence of the cognitive strategies used to generate training data. This design allows us to directly assess which generative strategies support effective latent variable extraction and generalization to human test data.

Each network was trained for 400 epochs with a batch size of 128 and a learning ate of 3 × 10−4, as outlined by Pan et al.^19^. To prevent overfitting, early stopping was applied after 35 epochs based on the loss calculated from the validation data. All networks converged to low training and validation losses (see Code Repository), indicating successful learning across models.

### Inference

During the inference phase (Figure 1D, adapted from Pan et al.^19^, the trained LaseNet takes the observable experimental data as input to infer a sequence of unobservable latent variables. Discrete latent states were inferred by selecting the state with the highest softmax probability at each time point (i.e., argmax over output probabilities), without applying a classification threshold, yielding a most-likely sequence of attentional focus across trials for synthetic (Figure 3) and human held-out test data (Figure 5).

Our test set comprised both synthetic and human self-report datasets that were not used during network training. To generate synthetic test data, we simulated an additional 1,000 unseen (*Z, Y*) sequences of 720 trials from each cognitive model. Because these datasets were generated under known models with known ground-truth attentional states, they provided a stringent test of each network’s ability to recover attentional dynamics produced by its training cognitive model.

To generate human test data, we collected trial-by-trial feature-level attention reports (e.g., “square”) from 21 participants completing the task (6 games, 18 trials per game). In this case, ground-truth attentional states were available via self-report, but the underlying generative cognitive model was unknown, allowing us to evaluate which networks inductive biases best captured human attentional dynamics.

### Network Performance Evaluation

For each trained network, we computed feature- and dimension-level labeling accuracy by passing the test datasets through the network and extracting trial-by-trial predictions of the attended feature (synthetic test accuracy: Figure 3; human test accuracy: Figure 5). Feature-level accuracy was defined as the proportion of trials in which the predicted feature matched the ground-truth label (chance = 0.167). Dimension-level accuracy was defined as the proportion of trials in which the predicted feature fell within the same dimension as the ground-truth label (e.g., both shape features; chance = 0.50).

To systematically evaluate decoding accuracy on human test data across networks trained under different cognitive models while accounting for repeated measurements within participants, we fit a linear mixed-effects model with decoding accuracy as the dependent variable (Eq. 1). Model identity was included as a categorical fixed effect, with the RS model specified as the reference level. A random intercept and slope were included for each participant. This analysis tested whether networks labeled human self-reported attentional focus significantly better than the RS baseline, which assumes no attention-learning mechanism. The equation was as follows:

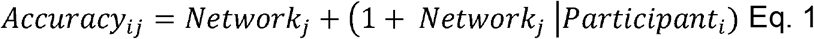

Where is *Accuracy*_*ij*_ denotes the decoding accuracy for participant i according to network *j. Network*_*j*_ represents a categorical fixed effect indexing the trained network (with RS specified as the reference level). The term (1 + *Network*_*j*_ ∣*Participant*_*i*_) indicates that both a random intercept and random slopes for Network were included for each participant, allowing baseline accuracy and the effect of Network on accuracy to vary across participants. Separate models were fit for attended feature- and dimension-labeling accuracy.

Following model fitting, we computed all pairwise contrasts among networks that incorporated attention-learning mechanisms (FRL, FRLd, Hybrid, rSHT, and PF). Pairwise contrasts were constructed as differences between the relevant fixed-effect estimates from the mixed-effects model. Standard errors for each contrast were derived from the fixed-effects covariance matrix, and Wald z-statistics were computed to assess significance. Two-sided p-values were obtained from the standard normal distribution. Because contrasts involving the RS baseline were directly tested within the primary mixed-effects model, they were excluded from post hoc comparison. Remaining pairwise p-values were corrected for multiple comparisons using the Benjamini– Hochberg FDR procedure.

### Network Recovery and Generalization Analyses

To assess network recoverability, we asked whether each trained network was most accurate when labeling synthetic test data generated from the cognitive model under which it was trained. Because synthetic datasets were generated under known cognitive models with known ground-truth attentional states, the correct network should, in principle, achieve the highest feature-label decoding accuracy on data produced by its corresponding generative model. Demonstrating this pattern would indicate that networks capture model-specific attentional dynamics rather than functioning as general-purpose decoders.

To quantify recoverability, we selected a subset of synthetic test data (100 simulated sequences per model; 720 trials per sequence) and evaluated each sequence using all trained networks. For each sequence, we identified the “best” network as the one achieving the highest feature-level decoding accuracy. We then computed the proportion of sequences for which the network corresponding to the generative model was the top-performing network (Supplementary Figure 3). This analysis treated performance categorically (best vs. not best) and did not consider graded differences in accuracy across networks. Sequences in which multiple networks achieved identical top accuracy were excluded (5 sequences in the rSHT test set and 5 in the Hybrid test set).

To further examine generalization, we focused on the FRLd and Hybrid networks and evaluated their feature-level decoding accuracy across all synthetic test datasets (1,000 sequences per generative model; Figure 7). Each network was applied to synthetic data generated from every cognitive model, allowing us to characterize the distribution of decoding performance across models. As expected, each network performed best on data generated from its own training model. Notably, performance was also elevated for models sharing mechanistic components within the same model class, indicating that networks generalized beyond their training distribution to related attentional dynamics rather than merely memorizing model-specific trajectories.

## Data and Code Availability

All data and analysis code necessary to reproduce the findings reported in this manuscript will be made publicly available in an open-source repository upon acceptance.

## References

1. Niv, Y. Learning task-state representations. Nat. Neurosci. 22, 1544–1553 (2019).

2. Radulescu, A., Shin, Y. S. & Niv, Y. Human Representation Learning. Annu. Rev. Neurosci. 44, 253–273 (2021).

3. Leukos, Mingze L., Liang, Albert, & Lindsay, Grace W. Modulation of feature attention by reward prediction error explains value learning behavior. Preprint at 10.64898/2026.04.10.717847 (2026).

4. Leong, Y. C., Radulescu, A., Daniel, R., DeWoskin, V. & Niv, Y. Dynamic Interaction between Reinforcement Learning and Attention in Multidimensional Environments. Neuron 93, 451–463 (2017).

5. Wilson, R. C. & Niv, Y. Inferring Relevance in a Changing World. Front. Hum. Neurosci. 5, (2012).

6. Song, M., Baah, P. A., Cai, M. B. & Niv, Y. Humans combine value learning and hypothesis testing strategically in multi-dimensional probabilistic reward learning. PLOS Comput. Biol. 18, e1010699 (2022).

7. Radulescu, A., Niv, Y. & Daw, N. A particle filtering account of selective attention during learning. in 2019 Conference on Cognitive Computational Neuroscience (Cognitive Computational Neuroscience, Berlin, Germany, 2019). doi:10.32470/CCN.2019.1338-0.

8. Niv, Y. et al. Reinforcement Learning in Multidimensional Environments Relies on Attention Mechanisms. J. Neurosci. 35, 8145–8157 (2015).

9. Radulescu, A., Daniel, R. & Niv, Y. The effects of aging on the interaction between reinforcement learning and attention. Psychol. Aging 31, 747–757 (2016).

10. Daniel, R., Radulescu, A. & Niv, Y. Intact Reinforcement Learning But Impaired Attentional Control During Multidimensional Probabilistic Learning in Older Adults. J. Neurosci. 40, 1084–1096 (2020).

11. Kruschke, J. K. ALCOVE: An Exemplar-Based Connectionist Model of Category Learning.

12. Roelfsema, P. R. & Ooyen, A. V. Attention-Gated Reinforcement Learning of Internal Representations for Classification. Neural Comput. 17, 2176–2214 (2005).

13. Bramlage, L. & Cortese, A. Generalized attention-weighted reinforcement learning. Neural Netw. 145, 10–21 (2022).

14. Mackintosh, N. J. SELECTIVE ATTENTION IN ANIMAL DISCRIMINATION LEARNING.

15. Jones, M. Integrating Reinforcement Learning with Models of Representation Learning.

16. Radulescu, A., Niv, Y. & Ballard, I. Holistic Reinforcement Learning: The Role of Structure and Attention. Trends Cogn. Sci. 23, 278–292 (2019).

17. Blakeman, S. & Mareschal, D. Selective particle attention: Rapidly and flexibly selecting features for deep reinforcement learning. Neural Netw. 150, 408–421 (2022).

18. Daw, N. D. & Courville, A. C. The pigeon as particle filter.

19. Pan, T.-F., Li, J.-J., Thompson, B. & Collins, A. Latent Variable Sequence Identification for Cognitive Models with Neural Network Estimators. Behav. Res. Methods 57, 272 (2025).

20. Chan, S. C. Y., Santoro, A., Lampinen, A. K., Wang, J. X. & Singh, A. K. Data Distributional Properties Drive Emergent In-Context Learning in Transformers.

21. Niv, Y. Learning task-state representations. Nat. Neurosci. 22, 1544–1553 (2019).

22. Bellman, R. Dynamic Programming. (Princeton Univ. Pr, Princeton, NJ, 1984).

23. Wise, T., Emery, K. & Radulescu, A. Naturalistic reinforcement learning. Trends Cogn. Sci. 28, 144–158 (2024).

24. Van Opheusden, B., Acerbi, L. & Ma, W. J. Unbiased and efficient log-likelihood estimation with inverse binomial sampling. PLOS Comput. Biol. 16, e1008483 (2020).

25. Wagner, AG, (second) & Rescorla, RA, (first). A theory of Pavlovian conditioning: variations in the effectiveness of reinforcement and nonreinforcement. in Classical conditioning II: current research and theory (Black AH, Prokasy WE eds) 64–99 (Appleton-Century-Crofts, New York).

26. Wilson, R. C. & Collins, A. G. Ten simple rules for the computational modeling of behavioral data. eLife 8, e49547 (2019).

