## Supplementary Information for "Learning to Attend Through Value-Based Hypothesis Testing"

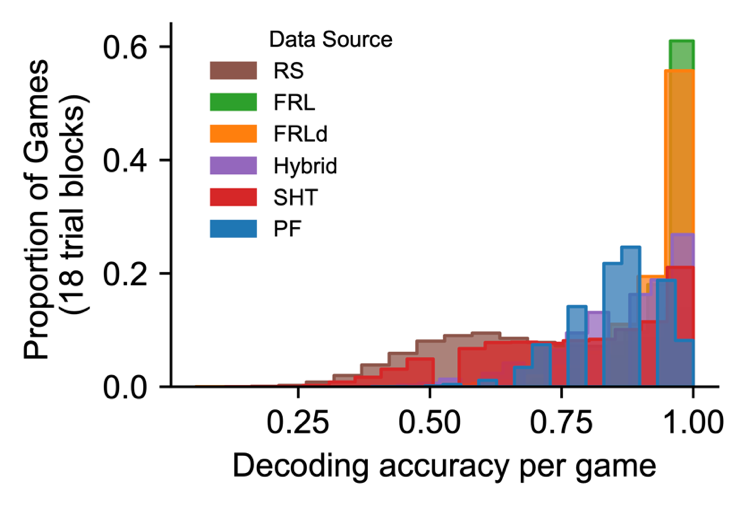

**Supplementary Figure 1. Distribution of decoding accuracy per game across networks**. Distribution of games (18 trials/game) that were perfectly labeled by each network. Single-game accuracy analyses revealed substantial variability across models in the proportion of games achieving 100% prediction accuracy (RS = 0.14, FRL = 0.61, FLRd = 0.56, Hybrid = 0.27, rSHT = 0.21, PF = 0.08). These results indicate that networks varied in their ability to fully recover trial-by-trial hypotheses within individual games; however, each network achieved perfect decoding in at least a subset of games.

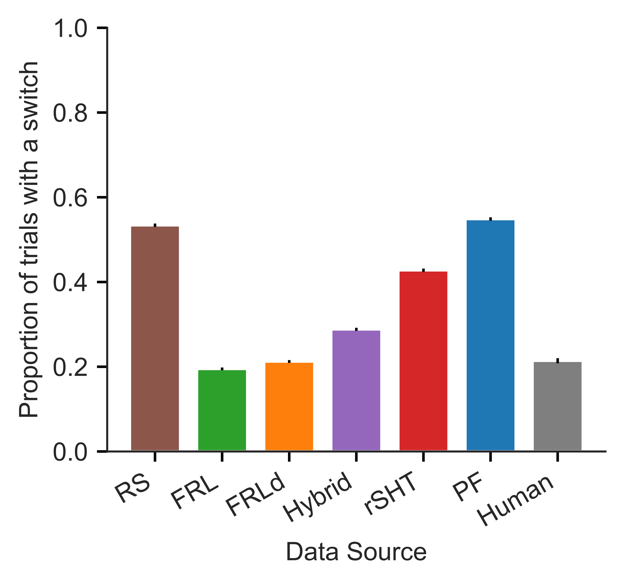

**Supplementary Figure 2. Attention switch rates across models and human data.**
Mean proportion of trials on which the hypothesized attended feature changed for each synthetic model and for human self-report data. For comparability, 21 synthetic agents were sampled from each model to match the number of human participants (N = 21 per group; mean switch rates: RS = 0.53, FRL = 0.19, FRLd = 0.21, Hybrid = 0.29, rSHT = 0.43, PF = 0.55, and humans = 0.21; error bars = SEM). While fast-switching mechanisms characteristic of SHT-based models produce high switch rates, human behavior is more conservative. The Hybrid model exhibits an intermediate switch rate, balancing excessive and insufficient switching across model classes.

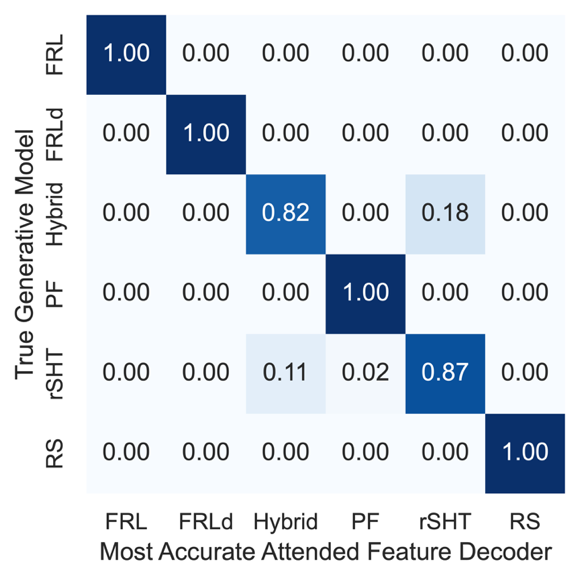

**Supplementary Figure 3. Network recovery across generative models.**
Each panel shows the proportion of simulations for which a given model’s network (columns) achieved the highest accuracy in decoding the attended feature for synthetic evaluation data generated by each model (rows; N = 100 simulated agents/model). Diagonal entries indicate correct model recovery, whereas off-diagonal entries reflect misidentification. Networks trained on the PF, FRL, FRLd, and RS models achieved perfect recovery (100%). The Hybrid and its component rSHT model were recovered with high but non-perfect accuracy (83% and 82%, respectively), with misclassifications occurring primarily between these two models. This overlap is expected, as under regimes of weak value learning the Hybrid model can express behavior similar to that of the SHT model. Overall, these results indicate that the networks learn largely dissociable behavioral strategies.
